# Heterogenous impairment of α-cell function in type 2 diabetes is linked to cell maturation state

**DOI:** 10.1101/2021.04.08.435504

**Authors:** Xiao-Qing Dai, Joan Camunas-Soler, Linford JB Briant, Theodore dos Santos, Aliya F Spigelman, Emily M. Walker, Rafael Arrojo e Drigo, Austin Bautista, Robert C. Jones, James Lyon, Aifang Nie, Nancy Smith, Jocelyn E Manning Fox, Seung K Kim, Patrik Rorsman, Roland W Stein, Stephen R Quake, Patrick E MacDonald

## Abstract

In diabetes, glucagon secretion from pancreatic α-cells is dysregulated. We examined α-cells from human donors and mice using combined electrophysiological, transcriptomic, and computational approaches. Rising glucose suppresses α-cell exocytosis by reducing P/Q-type Ca^2+^ channel activity, and this is disrupted in type 2 diabetes (T2D). Upon high-fat-feeding of mice, α-cells shift towards a ‘β-cell-like’ electrophysiologic profile in concert with an up-regulation of the β-cell Na^+^ channel isoform *Scn9a* and indications of impaired α-cell identity. In human α-cells we identify links between cell membrane properties and cell surface signalling receptors, mitochondrial respiratory complex assembly, and cell maturation. Cell type classification using machine learning of electrophysiology data demonstrates a heterogenous loss of ‘electrophysiologic identity’ in α-cells from donors with T2D. Indeed, a sub-set of α-cells with impaired exocytosis is defined by an enrichment in progenitor markers suggesting important links between α-cell maturation state and dysfunction in T2D.

**Key findings:**

- α-cell exocytosis is suppressed by glucose-dependent inhibition of P/Q-type Ca^2+^ currents
- Dysfunction of α-cells in type 2 diabetes is associated with a ‘β-cell-like’ electrophysiologic signature
- Patch-seq links maturation state, the mitochondrial respiratory chain, and cell surface receptor expression to α-cell function
- α-cell dysfunction occurs preferentially in cells enriched in endocrine lineage markers

## Introduction

In concert with reduced insulin secretion from pancreatic islet β-cells in type 2 diabetes (T2D), disrupted glucagon secretion from islet α-cells contributes to hyperglycemia and impaired hypoglycemia counter-regulation (Girard, 2017). While insulin and glucagon secretion are both dependent upon electrical excitability and Ca^2+^-dependent exocytosis, the complement and role of ion channels and the impact of glucose-stimulation differ in these cell types. Insulin granule exocytosis is linked to Ca^2+^ entry via L-type Ca^2+^ channels, whereas glucagon secretion is coupled to the P/Q-type Ca^2+^ channels (Marinis et al., 2010). Also, Na^+^ channels play a more prominent role in glucagon secretion (Barg et al., 2000; Göpel et al., 2000; Ramracheya et al., 2010) and differ in their regulation between endocrine cell types (Zhang et al., 2014). In rodents these differences can be used to distinguish cell types, either by the inactivation properties of Na^+^ currents (Zhang et al., 2014) or by the modelling of ‘electrophysiological fingerprints’ (Briant et al., 2017).

Similar to β-cells the excitatory and secretory machinery in α-cells, including ion channel activities (Huang et al., 2011a), intracellular Ca^2+^ responses (Marchand and Piston, 2010; Reissaus and Piston, 2017; Shuai et al., 2016), Zn^2+^ and glucagon content (Zadeh et al., 2020), and exocytotic capacity (Huang et al., 2011b) are heterogeneous. Indeed, the intracellular Ca^2+^ response of α-cells varies, with some suppressed by glucose and others activated (Shuai et al., 2016). This suggests a heterogeneity that is supported by single-cell transcriptomic findings (Camunas-Soler et al., 2020; Korsunsky et al., 2018).

Recent reports suggest a sub-population of α-cells (or at least ‘α-like-cells’) that are highly proliferative. In mice these are identified by up-regulation of the *Slc38a5* amino acid transporter (Kim et al., 2017). In humans these may be identified by the presence of ARX and cytosolic Sox9 (Lam et al., 2018). These cells could account for α-cell hyperplasia upon glucagon-receptor antagonism and may be a source of new β-cells via trans-differentiation (Meulen et al., 2017).

While little evidence so far exists to suggest that α-cells dedifferentiate, they have been shown to trans-differentiate in rodents following severe β-cell loss (Thorel et al., 2010) or genetic manipulation of transcription factor expression (Chakravarthy et al., 2017; Matsuoka et al., 2017). Interestingly, α-cells may show more plasticity than β-cells (Bramswig et al., 2013): they appear to exist in distinct states characterized by chromatin accessibility at promotors for *GCG*, functional genes such as *ABCC8*, and at sites enriched in motifs for binding of transcription factors for the RFX, GATA and NEUROD families, among others (Chiou et al., 2021). Human α-cells also appear to exhibit persistent immature transcript profiles which may contribute to impaired identity in T2D (Avrahami et al., 2020).

We hypothesised that this plasticity in α-cells influences their membrane function, contributing to α-cell dysfunction in T2D. We used correlated electrophysiological and single-cell RNAseq phenotyping (patch-seq) of α-cells from human donors and mice to identify a novel cell-autonomous glucose-regulation of α-cell Ca^2+^ channel activity and exocytosis that is associated with α-cell maturation state. Further investigation of transcriptional profiles that correlate with α-cell electrical function identify putative regulators of glucagon secretion, including the mitochondrial respiratory chain complex and numerous cell surface receptors. These results are complemented by studies where α-cells from mice fed a high fat diet adopt a ‘β-cell-like’ electrical profile, together with evidence for impaired α-cell identity. Similarly, in human T2D, α-cells enriched for markers of mitochondrial function and transcription factors such as *NEUROD1, ISL1, NKX2-2*, and *FEV* that define pancreatic endocrine lineage exhibit an impaired electrophysiological phenotype and dysregulated exocytosis. This suggests an important link between α-cell maturation state and α-cell dysfunction in T2D.

## Results

### Glucose-mediated suppression of α-cell exocytosis is disrupted in type 2 diabetes

In insulin-secreting β-cells, glucose metabolism amplifies Ca^2+^-triggered exocytosis and insulin secretion (Ferdaoussi et al., 2015; Gembal et al., 1992; Sato et al., 1992) which is linked to the activation of L-type Ca^2+^ channels (Barg et al., 2001; Bokvist et al., 1995; Wiser et al., 1999). We examined the impact of glucose on glucagon exocytosis in α-cells from donors with no diabetes (ND) or with T2D (**Suppl Table 1**) following dispersion to single cells and subsequent identification by glucagon immunostaining (**Fig 1A**). In ND α-cells, exocytosis was high at low glucose (1 mM) and increasing glucose (to 5-20 mM) suppresses this response. In α-cells from donors with T2D, exocytosis was ∼75% lower at 1 mM glucose and increasing glucose exerted no additional inhibitory effect (**Fig 1B**; **Suppl Fig 1**). These differences are also obvious when cells are grouped by individual donor (**Fig 1C**) and are consistent with a recent study examining glucagon exocytosis by live cell imaging (Omar-Hmeadi et al., 2020). In concert, glucose suppresses the activation of voltage-dependent Ca^2+^ channel (VDCC) activity in ND α-cells (**Fig 1D**; **Suppl Fig 1**). The majority of the human α-cell Ca^2+^ current is mediated by L-type (∼30%) and P/Q-type (∼70%) channels (Ramracheya et al., 2010), and the latter are directly linked to glucagon exocytosis (Dai et al., 2014; Ramracheya et al., 2010; 2018). Accordingly, the P/Q-channel blocker agatoxin, but not the L-channel blocker isradipine, inhibits exocytosis from human ND α-cells at 1 mM glucose (**Fig 1E**). Finally, the glucose-regulated Ca^2+^ current in human ND α-cells is mediated by agatoxin-sensitive P/Q-type channels but is absent at 1 mM glucose in T2D α-cells (**Fig 1F**; **Suppl Fig 1**).

**Figure 1.**
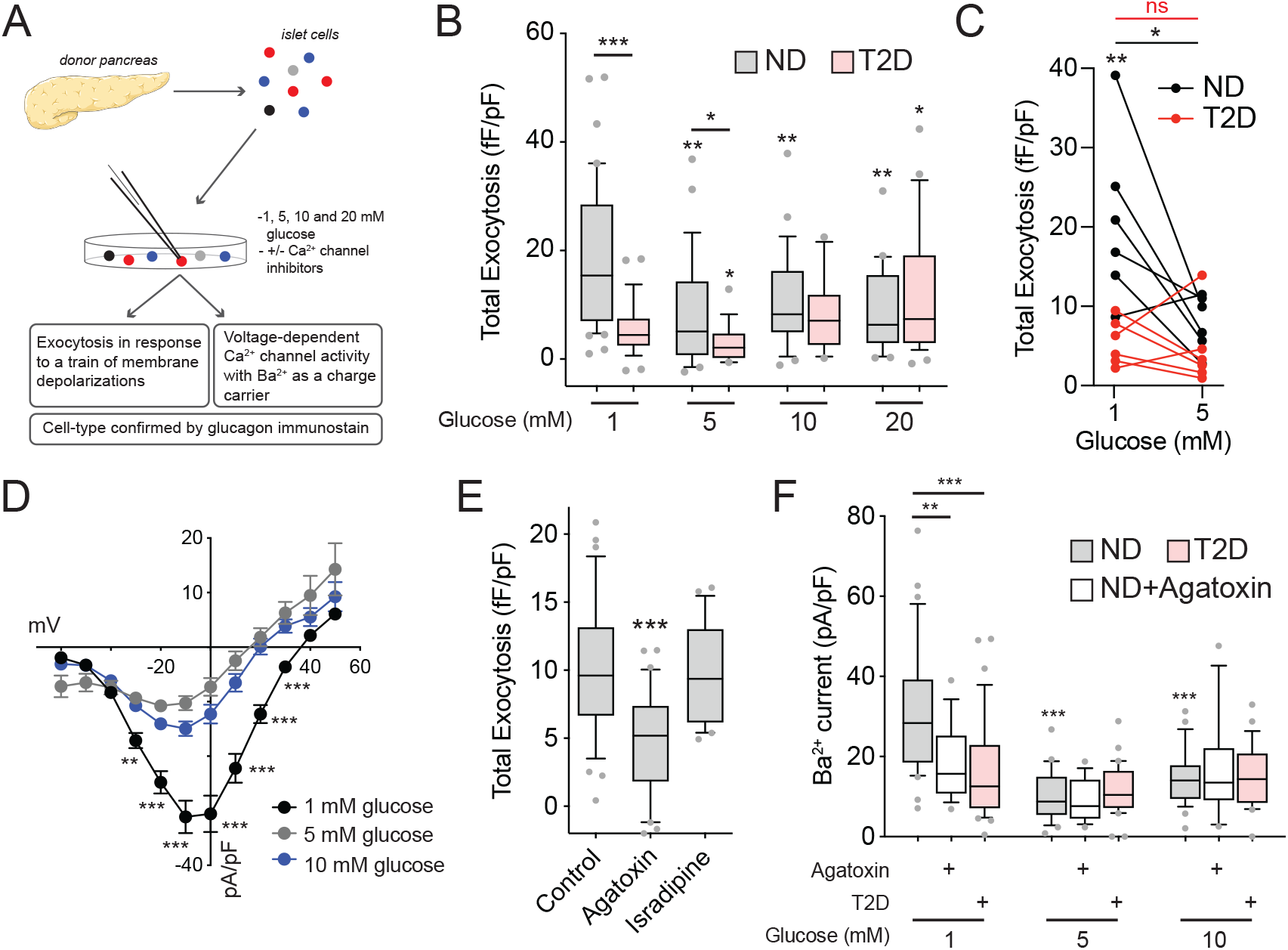
Glucose suppresses human α-cell exocytosis in concert with P/Q-channel activity. **A)** Human islets were isolated, dispersed and cultured for 1-2 days prior to electrophysiological recording of exocytosis or Ca^2+^ channel properties. The α-cells were subsequently identified by glucagon immunostaining. **B)** Total exocytosis in ND α-cells is suppressed by increasing glucose (*grey*, n=42, 27, 24, 20 cells from 9 donors), but this is disrupted in T2D α-cells (*pink*, n=31, 20, 13, 28 cells from 6 donors). **C)** Averaged values from B shown for individual donors (n=6 ND, 6 T2D) with paired data at 1 mM and 5 mM glucose. **D)** Ca^2+^ channel activity in ND α-cells is suppressed by increasing glucose (n=31, 24, 27 cells from 7 donors). **E)** In ND α-cells exocytosis (at 1 mM glucose) is suppressed by the P/Q-type Ca^2+^ channel blocker agatoxin (100 nM), but not the L-type Ca^2+^ channel blocker isradipine (10 μM; n=34, 30, 21 cells from 5 donors). **F)** Ca^2+^ channel activity (measured using Ba^2+^ as a charge carrier) is suppressed by agatoxin (100 nM) in ND α-cells to the level seen in T2D α-cells (n=31, 16, 24, 15, 27, 18 cells from 7 ND donors, and 34, 31, 27 cells from 5 T2D donors). Data were compared by one-way or two-way ANOVA, followed by Tukey post-test to compare groups or the Student’s t-test (panel C). *-p<0.05; **-p<0.01; and ***-p<0.001 compared with 1 mM glucose control, versus the ND control (panel C) or as indicated.

Consistent with an acute impact of glucose on depolarization-induced glucagon exocytosis in human cells, glucagon secretion from mouse islets was increased by switching from 5 to 1 mM glucose (**Fig 2A**). Following an ∼10-minute exposure to 1 mM glucose (*black line*, **Fig 2A**), direct depolarization with 20 mM KCl elicited a robust but transient stimulation of glucagon release that was blunted in islets kept at 5 mM glucose (*blue line*, **Fig 2A**). These differences could not be explained by the indirect paracrine effects of insulin, and insulin secretion evoked by high K^+^ stimulation was the same whether at 1 or 5 mM glucose (**Fig 2B**). Similar to human α-cells, increasing glucose suppresses exocytosis in mouse α-cells (**Fig 2C**). Although the physiological impact of glucose on glucagon secretion involves key paracrine signals (Briant et al., 2016), the suppression of exocytosis produced by 5 mM glucose was similar when cells were seeded at a cell density that was 10% of normal to minimize the potential impact of paracrine signaling (**Fig 2D**) and requires glucose metabolism since the non-metabolizable glucose analog 2-deoxyglucose (2-DG) does not mimic the effect of glucose (**Fig 2E**).

**Figure 2.**
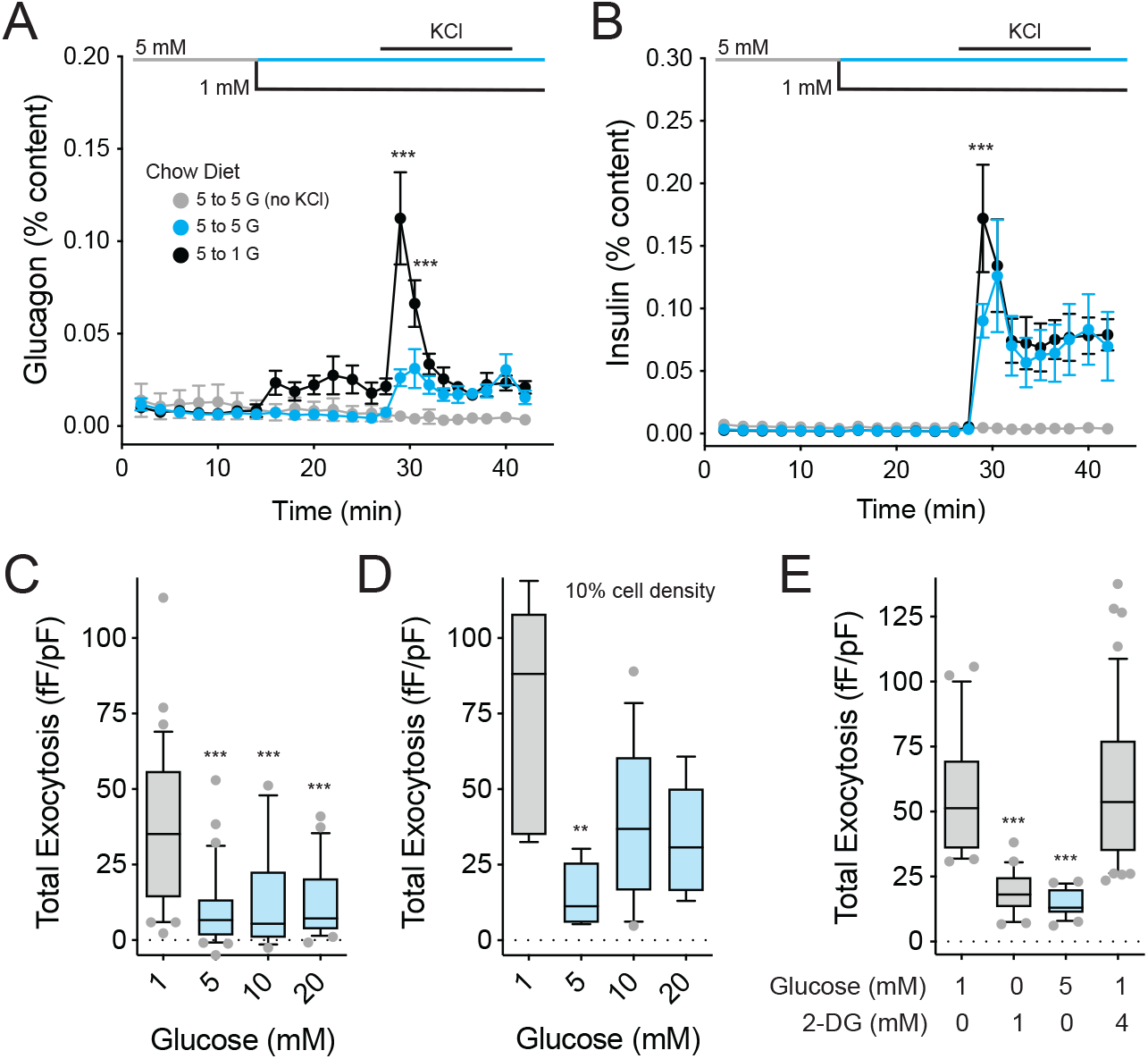
Glucose-suppression of mouse α-cell exocytosis requires glucose metabolism. **A-B)** Glucagon secretion (A) from islets of male C57bl6 mice increases as glucose drops from 5 mM to 1 mM glucose (*black*). The glucagon response to direct depolarization of islets with 20 mM KCl is much greater at 1 mM (black) than at 5 mM (*blue*) glucose. Changes in insulin secretion (B) cannot explain this. A control maintained at 5 mM glucose is also shown (*grey*). (n=3, 3, 3 mice) **C-D)** Mouse α-cell exocytosis is suppressed by glucose (n=32, 30, 14, 27 cells from 4 mice), and this is preserved when cells are seeded at very low (10% of normal) density (D; n=6, 6, 13, 9 cells from 2 mice). **E)** Minimal glucose metabolism is required to support α-cell exocytosis, and also to suppress exocytosis when elevated as shown using the non-metabolizable glucose analog 2-Deoxy-D-glucose (2-DG) (n= 21, 20, 21, 46 cells from 4 mice). Data were compared by one-way or two-way ANOVA, followed by Tukey post-test to compare points or groups. **-p<0.01; and ***-p<0.001 compared with the 5 mM glucose points (panels A and B) or with the 1 mM glucose control group.

### Patch-seq highlights the role of mitochondrial respiratory chain in α-cell exocytosis, and suggests poor responsiveness in immature α-cells

We measured depolarization-induced exocytosis in concert with single-cell RNA-seq (patch-seq) and compared the responses of 400 α-cells at 1 and 10 mM glucose (31 donors; **Fig 3A**). In these cells, we find a glucose-dependent shift in the distribution of exocytosis-transcript correlations consistent with the suppressive effect of glucose on α-cell exocytosis (**Fig 3B**; **Suppl Table 2**). In other words, we observed an enrichment of positively-correlated genes to α-cell exocytosis at 1 mM (where exocytosis is high) and an enrichment of negatively-correlated genes at 10 mM glucose (where exocytosis is suppressed). Gene-set enrichment analysis (GSEA) using Z-scores of transcripts found in at least 20% of cells highlights pathways associated with elevated α-cell exocytosis at 1 mM glucose and suppression of exocytosis at 10 mM glucose (**Fig 3C**). Consistent with a role for metabolism in determining α-cell responsiveness, we find transcripts associated with mitochondrial respiratory chain complex assembly as positive correlates to exocytosis at low glucose and as negative correlates when glucose increases (**Fig 3C-D**). A separate over-representation analysis (ORA) of significant transcriptional correlates yields similar results and highlights electron transport, ATP synthesis, protein synthesis and protein targeting as correlates of robust α-cell exocytosis at 1 mM and suppression at 10 mM glucose (**Suppl Fig 2**).

**Figure 3.**
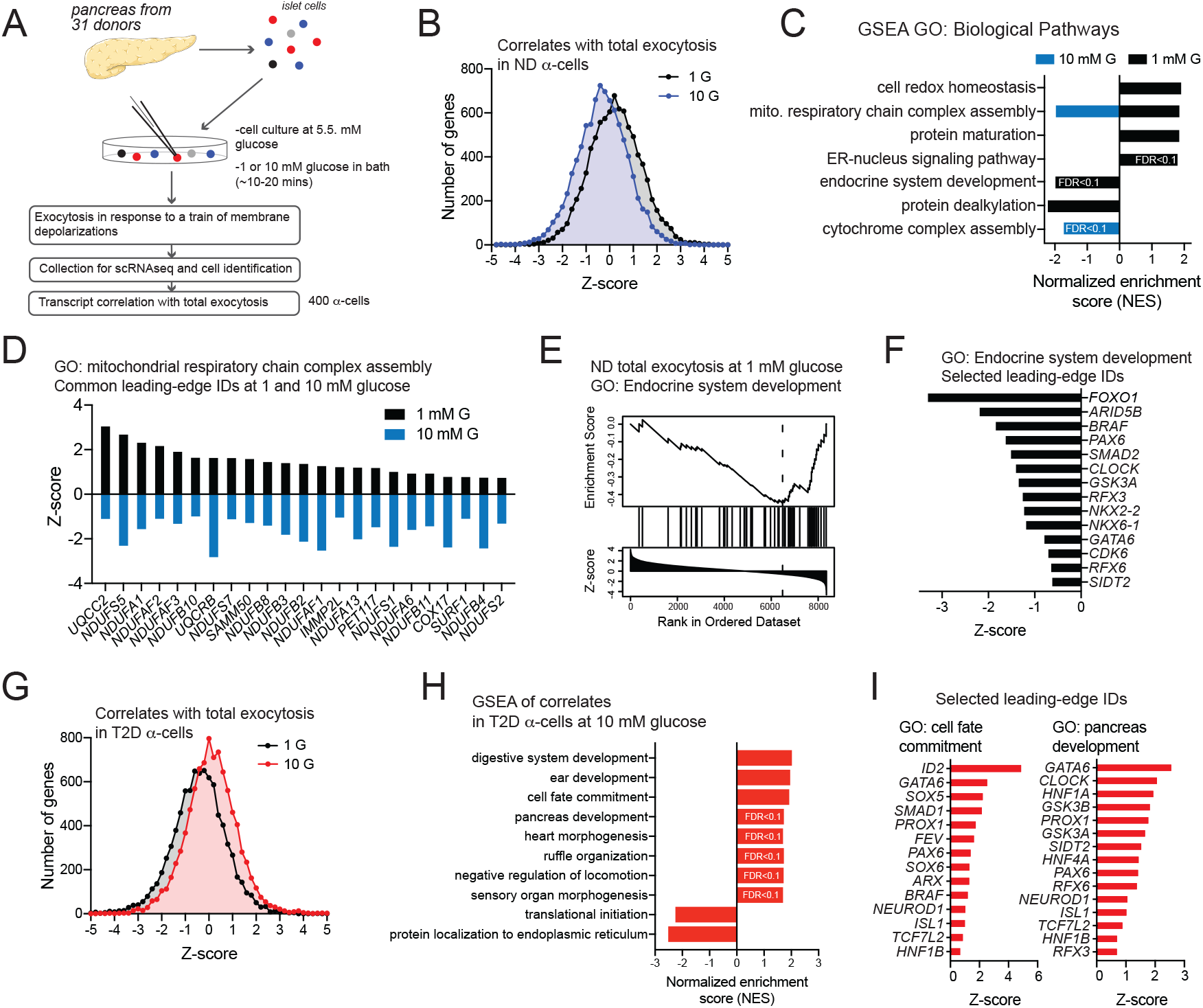
Impaired α-cell exocytosis in T2D is linked to endocrine development and cell fate. **A)** Donor islet cells were assessed by patch-clamp electrophysiology and collected for scRNAseq. 400 α-cells from 24 ND and 7 T2D donors were assessed by patch-seq. These some cells for which data have been partly published previously (Camunas-Soler et al., 2020). **B)** The distribution of ND α-cell exocytosis - transcript correlations shift leftward as glucose increases, consistent with suppression of exocytosis at 10 mM glucose. **C)** Gene set enrichment analysis (GSEA) using Z-scores as weighting across the entire transcriptome (using genes expressed in at least 20% of α-cells) reveals pathways linked to either facilitation of exocytosis (positive values) or suppression of exocytosis (negative values). FDR<0.05 unless indicated otherwise. **D)** Mitochondrial respiratory chain complex assembly is associated both with robust α-cell exocytosis at 1 mM glucose and suppression of exocytosis at 10 mM glucose, and the common leading-edge transcripts that underly this pathway flip their correlation as glucose increases. **E)** An endocrine system development pathway appears enriched in ND α-cells that fail to show robust exocytotic responses at 1 mM glucose. **F)** Several leading-edge IDs underlying the signal in panel E include transcripts involved in islet cell lineage and α-cell identity. **G)** In T2D α-cells the distribution of transcript correlates to exocytosis shift rightward as glucose increases, consistent with an inappropriately elevated exocytosis at 10 mM glucose. **H)** GSEA with Z-scores as weighting (using genes expressed in at least 20% of α-cells) reveals several development and cell fate pathways that appear linked to inappropriate increases in T2D α-cell exocytosis at 10 mM glucose (FDR<0.05 unless indicated otherwise). **I)** Selected leading-edge transcripts that underlie the cell fate and pancreas development pathways highlight several islet lineage and α-cell identity markers. False discovery rate (FDR) for pathways identified by GSEA were <0.05 unless indicated otherwise.

Transcripts associated with endocrine development appear as anti-correlates of exocytosis at 1 mM glucose (**Fig 3E-F**), suggesting a role for cell differentiation state as a determinant of robust responsiveness of α-cells in ND donors. In T2D α-cells, the effect of glucose on the distribution of transcript correlations is reversed, such that increasing glucose now associates with more positively correlated genes (**Fig 3G**; **Suppl Table 3**), consistent with our observation of impaired glucose-regulation of α-cell exocytosis in T2D. GSEA of these correlations highlights a role for cell development state in the inappropriately high α-cell exocytosis at 10 mM glucose in T2D (**Fig 3H**). Several leading-edge transcripts enriched in these pathways, including transcription factors important for pancreatic endocrine maturity like *GATA6, PAX6, RFX6*, and *RFX3*, overlap with those that correlate with inappropriately low exocytosis in ND α-cells at 1 mM glucose (**Fig 3F**,**I**). This raises the possibility that impaired glucose-regulation of α-cell exocytosis is related to developmental state or impaired identity (Avrahami et al., 2020).

### High fat diet impairs the ‘electrophysiological identity’ of mouse α-cells

Lowering glucose from 5 to 1 mM stimulates glucagon secretion by 2-fold. This was enhanced in mice fed a high fat diet (HFD) for 12-14 weeks (**Fig 4A**), similar to what we have shown previously (Kellard et al., 2020). There was no measurable difference in insulin secretion at 5 and 1 mM glucose but the response to high-K^+^ was reduced (**Fig 4B**), possibly reflecting the disruption of a direct coupling between Ca^2+^ channels to the insulin granules (Collins et al., 2010). As in human α-cells, exocytosis at 1 mM glucose in α-cells from chow fed mice is inhibited by the P/Q-type Ca^2+^ channel blocker agatoxin but not the L-type channel blocker isradipine (**Fig 4C**). This response is reversed in α-cells from HFD mice where exocytosis is resistant to agatoxin but sensitive to isradipine (**Fig 4C**). Indeed, the P/Q-channel agonist GV-58 (Tarr et al., 2012), which acts by delaying channel inactivation (**Fig 4D**), can ‘rescue’ exocytosis following suppression by 5 mM glucose in α-cells from chow-fed, but not HFD-fed, mice consistent with a loss of P/Q- channel coupling to exocytosis (**Fig 4E**). Mouse α- and β-cells are distinguished by a characteristic right-shifted Na^+^ current half-inactivation in α-cells compared with β-cells, which primarily express *Scn3a* and *Scn9a*, respectively (Zhang et al., 2014). Many HFD α-cells, identified by glucagon immunostaining, show a leftward shift in Na^+^ channel half-inactivation towards a ‘β-cell-like’ half-inactivation (**Fig 4F**).

**Figure 4.**
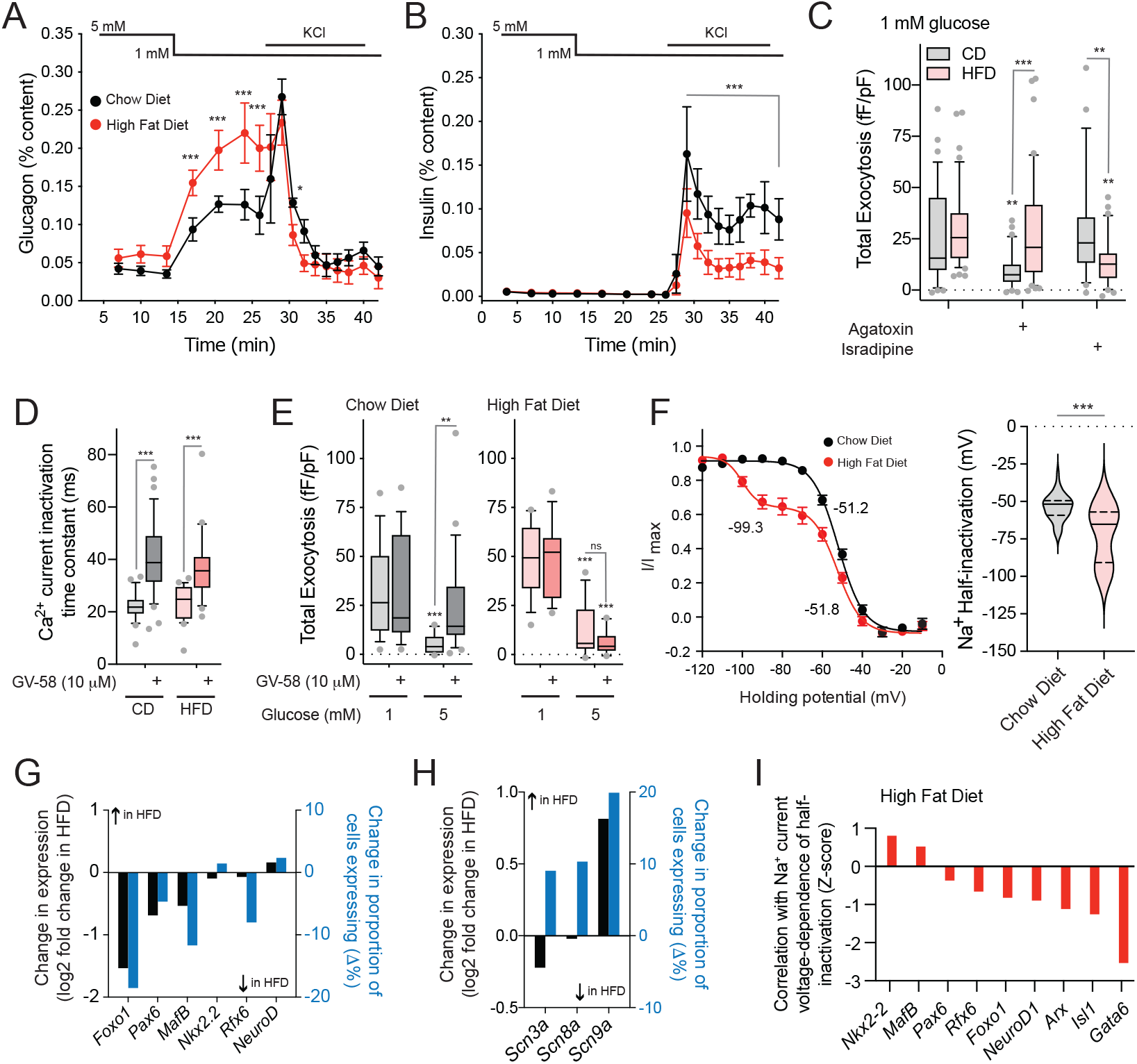
‘β-cell like’ properties of α-cells from high fat fed mice. **A-B)** Hypersecretion of glucagon (A) from islets of male HFD mice (*red*) compared with age matched chow fed controls (*black*) cannot be explained by differences in insulin secretion (B). (n=3, 3 mice). **C)** At 1 mM glucose mouse α-cell exocytosis is dependent on P/Q-type Ca^2+^ channels rather than L-type channels (shown by blockers agatoxin and isradipine at 100 nM and 10 μM, respectively), but this is reversed in HFD α-cells. (n=34, 46, 34, 41, 31, 40 cells from 8 HFD and 8 CD mice). **D-E)** The P/Q-channel activator GV-58 increase Ca^2+^ currents in both CD and HFD α-cells (D, n=23, 36, 24, 23 cells from 3 CD and 3 HFD mice) and can rescue glucose-suppressed exocytosis (E) in CD α-cells (*grey*; n=15, 16, 16, 21 cells from 3 CD mice) but not HFD α-cells (*pink*, n=14, 13, 15, 14 cells from 3 HFD mice). **F)** The α-cells from HFD mice show a leftward shift in the voltage-dependent inactivation of Na^+^ currents. (n=36, 44 cells from 9 CD and 9 HFD mice, measured at 1 mM glucose). Half-inactivation values from fit curves are indicated. **G)** From a separate set of CD and HFD α-cells assessed by patch-seq (**Suppl Fig 3**), mean expression (*black*) and percentage of cells expressing (*blue*) some α-cell identity transcription factors decrease in HFD α-cells compared with CD controls. **H)** Mean expression (*black*) and percentage of cells expressing (*blue*) the β-cell Na^+^ channel isoform, *Scn9a*, increases in HFD α-cells. **I)** The negative shift in Na^+^ channel steady-state inactivation in HFD α-cells correlates with transcripts involved in α-cell linage and identity. Data were compared by one-way or two-way ANOVA, followed by Tukey post-test to compare points or groups, or the Student’s t-test (panel F). *-p<0.05; **-p<0.01; and ***-p<0.001 compared with the 5 mM glucose points (panels A and B), with the 1 mM glucose control group, or as indicated.

This switching of both Ca^2+^-channel-exocytosis coupling and Na^+^ channel half-inactivation upon HFD, towards ‘β-cell-like’ phenotypes, is reminiscent of our observations during genetically induced α-to β-cell trans-differentiation confirmed by lineage tracing (Chakravarthy et al., 2017). However, the present changes occur while glucagon expression is maintained since the α-cells were identified by glucagon immunostaining. We explored these findings in a separate series of experiments where α-cells from HFD mice were collected for patch-seq (**Suppl Fig 3**). We found that HFD feeding evoked a down-regulation of some endocrine transcription factors (**Fig 4G**; **Suppl Table 4**), an upregulation of the β-cell Na^+^ channel transcript *Scn9a* (**Fig 4H**), and a correlation between the HFD-induced shift in Na^+^ channel inactivation and expression of α-cell identity and islet lineage transcription factors (**Fig 4I**; **Suppl Table 5**). Thus, after high fat feeding a population of mouse α-cells adopt electrophysiological properties typically associated with β-cell function. The transcriptional profile of these cells indicate that this shift is associated with altered α-cell identity and/or maturation state.

### Electrophysiological fingerprint modelling links human α-cell behavior and cell phenotype

We next compiled data from islet cells of 67 donors collected at 1, 5 and 10 mM glucose, and identified either by scRNAseq or immunostaining (**Fig 5A**). Unlike what is seen in mice, but similar to previous reports in human cells (Braun et al., 2008; Ramracheya et al., 2010), Na^+^ channel inactivation properties are very similar human α- and β-cells, with the peak Na^+^ currents (**Fig 5B**) and voltage-dependencies of inactivation (**Fig 5C**) being broadly distributed and overlapping between cell types. In ND α-cells the peak Na^+^ current significantly correlated with genes known to impact α-cell activity (**Fig 5D**; **Suppl Table 6**). Top positive correlates include genes involved in α-cell or islet function; including Na^+^ and Ca^2+^ channels (*SCN3A, SCN3B, CACNA1A*, etc.), α- and islet-cell lineage transcription factors (*MAFB, FEV, NKX2*.*2, NEUROD1*), G-protein coupled receptors known to impact α-cell function (*GIPR, SSTR1, FFAR1, GPR119*), and secreted factors implicated in islet function or development (*LOXL4, WNT4, UCN3, IL16*). Significant negative correlates were less abundant but included the vitamin D binding protein (*GC*) which is known to regulate α-cell Na^+^ currents (Viloria et al., 2020). GSEA for KEGG Pathways and GO: Biological Process terms revealed additional pathways (FDR<0.05) enriched in ND α-cells that show larger Na^+^ currents (**Suppl Fig 4**). Neurotransmitter signaling, such as serotonin, GABA and glutamate pathways, are correlated to α-cell Na^+^ channel activity. Numerous cell surface receptor transcripts also appear as correlates to Na^+^ channel activity (**Fig 5E, Suppl Fig 4**). We validated the ability of agonists for some of these to increase peak human α-cell Na^+^ currents (**Fig 5F**).

**Figure 5.**
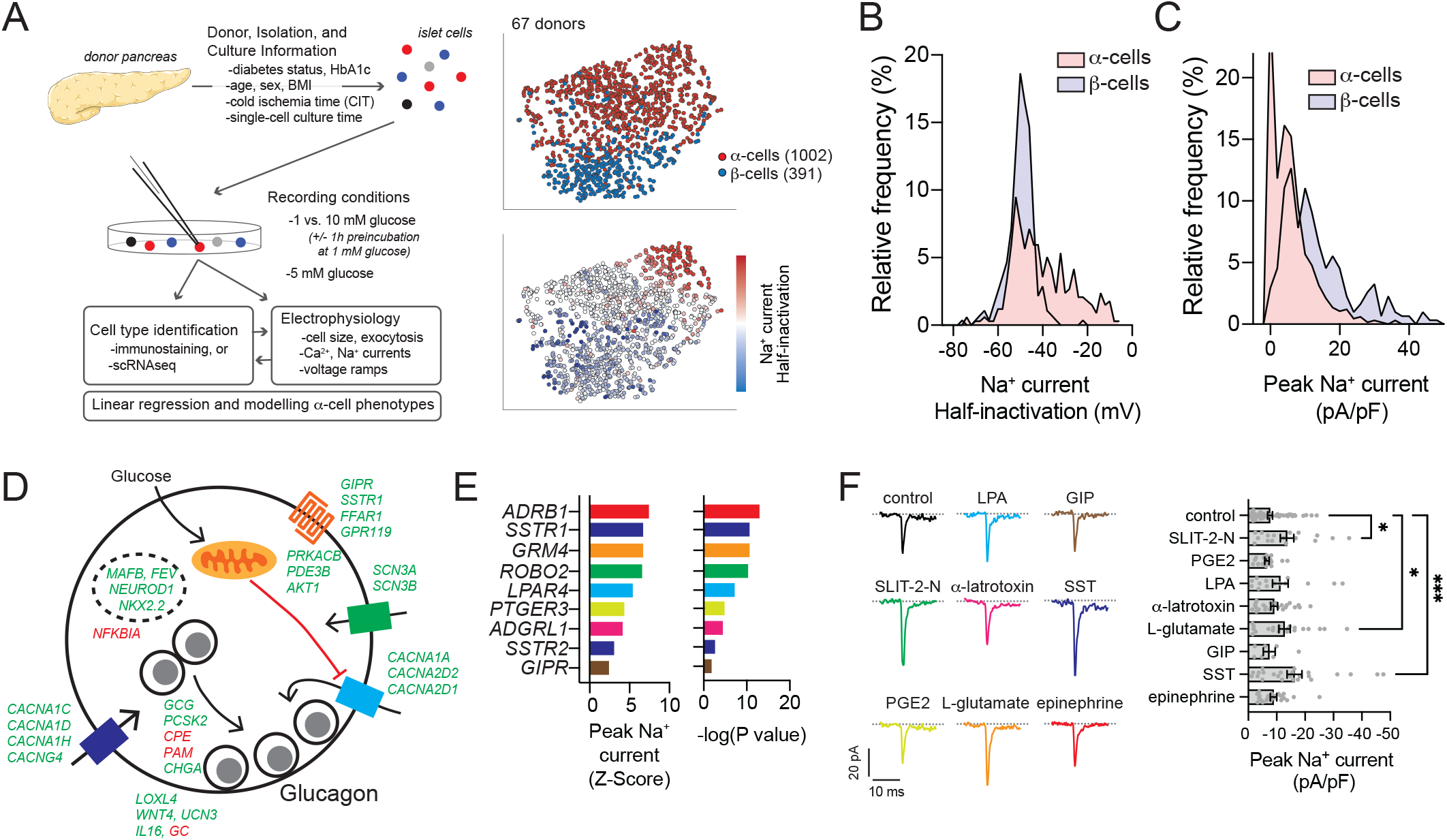
Na^+^ current properties correlate with transcriptomic markers of α-cell function. **A)** Outline of an expanded dataset of electrically profiled human islet cells, and tSNE representations of α- and β-cells identified by immunostaining or sequencing along with the relative distribution of Na^+^ current half-inactivation values. **B-C)** The distribution of Na^+^ current amplitudes (B) and voltage-dependence of half-inactivation values (C) demonstrate significant heterogeneity and overlap between β-cell (*light blue*) and α-cell (*pink*) populations. **D)** Correlation between α-cell peak Na^+^ current and transcript expression reveals several significant positive (*green*) and negative (*red*) correlations, some of which are mapped here (see also **Suppl Table 5**). **E)** Correlation of α-cell peak Na^+^ current and selected transmembrane signalling receptor transcripts (see also **Suppl Fig 4**). **F)** In separate donors, Na^+^ currents measured with receptor agonists (colours matching receptors shown in panel E) upon depolarization from -70 to -10 mV. Peak current is shown at right: control (n=53 cells); 0.5 μg/ml SLIT-2-N (n=17 cells); 10 μM prostaglandin E_2_ (PGE2, n=17 cells); 0.5 μM lysophosphatidic acid (LPA, n=14 cells); μg/ml α-latratoxin (n=24 cells); 10 mM L-glutamic acid (n=21 cells); 100 nM glucose-dependent insulinotropic polypeptide (GIP, n=7 cells); 200 nM somatostatin (SST, n=24 cells); 5 μM epinephrine (n=22 cells). Data were compared by one-way ANOVA, followed by the Benjamini and Hochburg post-test to method to compare groups controlling for false discovery rate. *-p<0.05 and ***-p<0.001 compared with control.

Although α-cell lineage markers correlate with Na^+^ current activity, the high degree of overlap in the Na^+^ current properties between human α- and β-cells make it impossible to use Na^+^ currents alone to interrogate shifts in human α-cell phenotypes. We therefore developed ‘electrophysiological fingerprinting’ classifier models (**Fig 6A**) that integrate Na^+^ current, Ca^2+^ current and exocytosis measures (**Fig 6B**) to characterize human α-cell ‘functional phenotypes’, similar to the logistic regression model (Briant et al., 2017) and algorithms based on ‘random forests’ (Camunas-Soler et al., 2020) we used previously. In those models, cell size was the major predictor of α-cell identity. To interrogate electrical phenotypes based on cell function, rather than cell size, we generated three independent models for cell type classification: Optimizable Ensemble classifiers that include (Model 1) or exclude (Model 2) cell size as an independent variable, and an Extreme Gradient Boosting model that also excludes cell size, but with additional restrictions to donor age, donor BMI and organ cold ischemic time (CIT) applied to the training data (Model 3). These properties distinguish ND α- and β-cells (**Fig 6A**) and, unlike some underlying electrophysiological parameters, were unaffected by potential confounders such as donor sex, BMI, CIT, cell culture time, and glucose concentration (**Fig 6B**).

**Figure 6.**
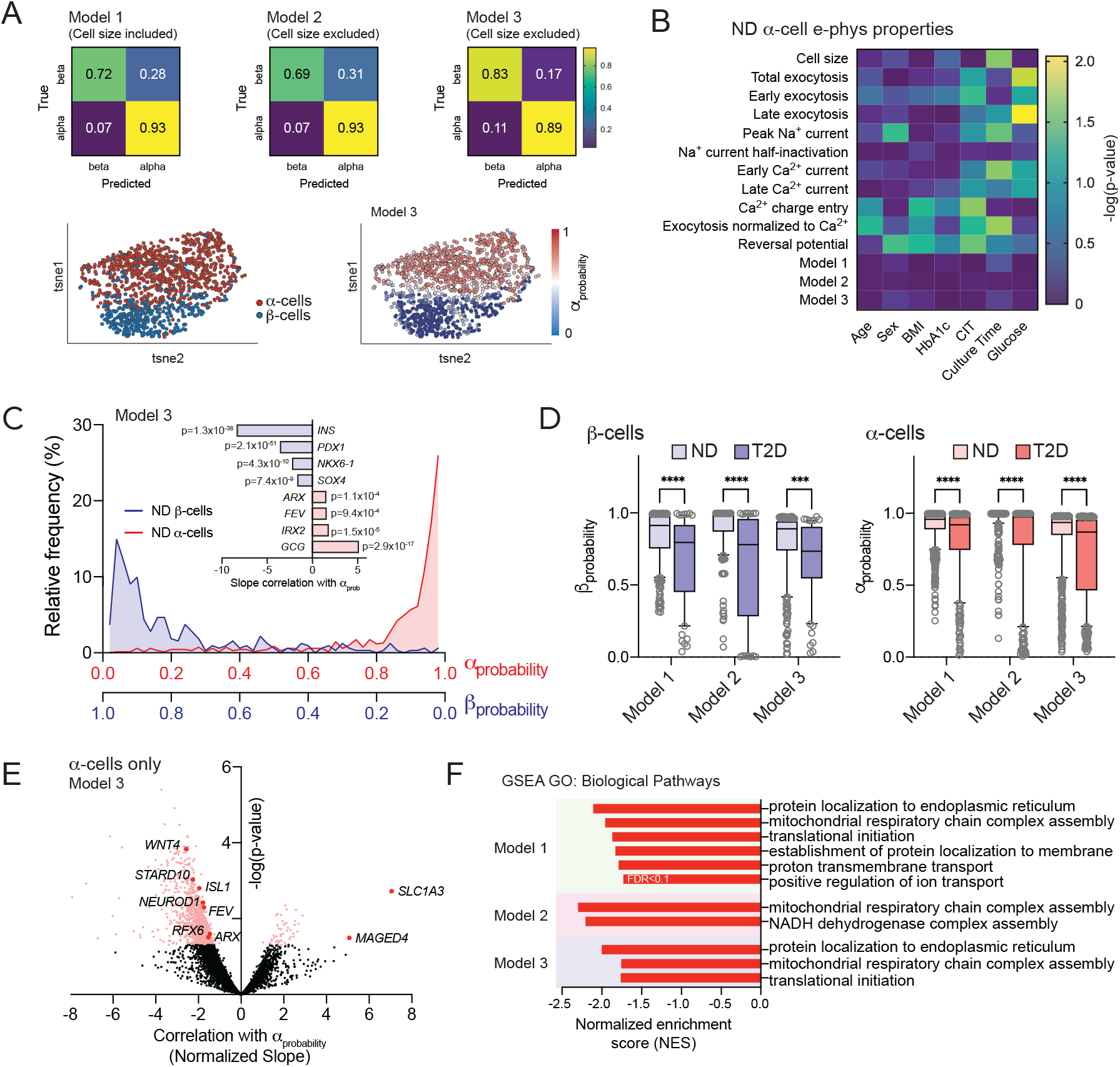
Electrophysiological fingerprints define a loss of ‘functional identity’ in T2D α-cells. **A)** Classifier models trained on islet cell electrical properties of α- and β-cells from donors with no diabetes (ND) using Optimizable Ensemble or Extreme Gradient Boosting (XGBoost) approaches, correctly identify cell types with high accuracy regardless of the inclusion or exclusion of cell size from training data. 80% of data was used for training. 20% of data was reserved for further validation. Assigned α_probability_ scores closely matched cell types determined by immunostaining or sequencing. **B)** Ordinary least squares multiple regression of ND α-cell electrophysiological properties, Model scoring, and donor/isolation variables. **C)** When applied to cells collected for patch-seq analysis, α_probability_ values correlate with high significance to the expression of canonical β-cell (*light blue*) and α-cell (*pink*) markers. **D)** In type 2 diabetes (T2D), both μ- and α-cells exhibit a ‘loss of electrophysiological phenotype’ indicated by a reduced fit to the three models. **E)** Volcano plot of transcript correlations with Model 3 α_probability_ values (slope/standard deviation) in T2D α-cells. **F)** Gene Set Enrichment Analysis (GSEA) reveal a role for pathways that include the mitochondrial respiratory chain complex in reduced α_probability_ of T2D α-cells in all three Models. Data in panel D were compared within models using non-parametric Kruskal-Wallis test followed by Dunn’s post-test to correct for multiple comparisons. ***-p<0.001 and ****p<0.0001 as indicated. False discovery rate (FDR) for pathways identified by GSEA were <0.05 unless indicated otherwise.

Model output between 0 and 1 can be considered the probability an electrophysiological profile matches that of an α-cell (1 = high confidence the cell is an α-cell) and we therefore called this α_probability_ (the converse being β_probability_). Assignment of scores to all ND α-and β-cells for which we had patch-seq data, without *a priori* knowledge of cell type, showed the effective separation of cell types and correlation with canonical markers (**Fig 6C**). All three models show a significantly reduced β_probability_ and α_probability_, in μ- and α-cells respectively, from donors with T2D (**Fig 6D**). The expression of numerous transcripts correlate with this ‘loss of electrophysiological identity’ in T2D α-cells within one or more of the models, including endocrine lineage markers such as *ISL1, NEUROD1, FEV*, and *RFX6* (**Fig 6E**; **Suppl Table 7**). GSEA using α_probability_ correlation slopes as weighting for transcripts expressed in at least 20% of cells highlight GO: Biological Process terms related to mitochondrial respiratory chain complex assembly in all three models (**Fig 6F**), similar to what we found for the glucose-regulation of α-cell exocytosis.

### Impaired functional identity and exocytosis in T2D α-cells enriched in lineage and identity markers

Human α-cells likely exist in distinct states characterized by chromatin accessibility at the *GCG* promoter and sites enriched with transcription factor motifs characterizing endocrine lineage and development (Chiou et al., 2021). Markers of α-cell lineage are heterogenous within our dataset, with evidence for increased expression of transcription factors involved in endocrine lineage specification in T2D (**Fig 7A**; **Suppl Fig 5**). Intriguingly, this also includes *ARX* which we confirm at the protein level, along with MAFB and glucagon itself (**Fig 7B**). *ARX*-enriched (*ARX*^hi^) cells express consistently higher levels of progenitor transcripts like *NEUROD1, FEV, GATA6*, and others (**Suppl Fig 5**). We find no difference in electrophysiological profiles (i.e. α_probability_ scores) between ND α-cells either enriched or deficient in these (**Fig 7C**; **Suppl Fig 5**). In T2D however, impaired electrical function occurs selectively in α-cells with higher levels of *ARX, NEUROD1, ISL1, PAX6, GATA6, FEV, NKX2-2, RFX6* and *MAFB* (**Fig 7C**; **Suppl Fig 5**). Accordingly, *ARX* and these other α-cell identity and linage transcripts do not correlate with exocytosis in ND α-cells, but correlate significantly with impaired exocytosis at 1 mM glucose in T2D (**Fig 7D**). In T2D, α-cells with low *ISL1, NKX2-2, NEUROD1* and *ARX* have normal exocytosis at low glucose, while T2D α-cells enriched in these show clearly impaired exocytotic function (**Fig 7E**).

**Figure 7.**
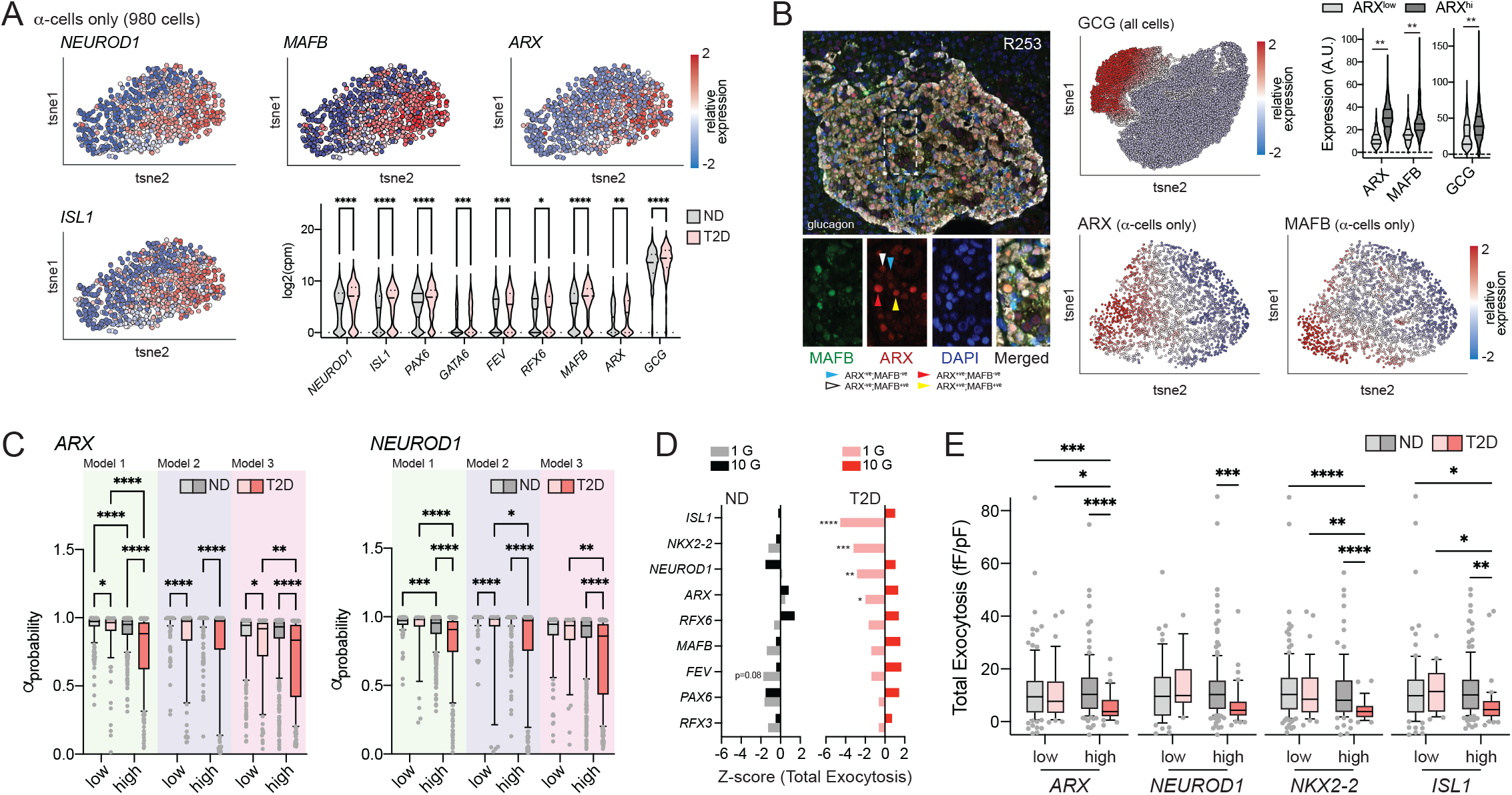
A role for maturation state in α-cell dysfunction in T2D. **A)** Heterogenous expression of transcript makers for islet cell lineage and α-cell maturity (980 patch-seq α-cells). **B)** We confirmed that ARX and MAFB are heterogeneously expressed in α-cells at the protein level *in situ* in the donors we have studied. Violin plots show the relative levels of GCG, ARX and MAFB protein expressed in ARX^low^ and ARX^hi^ α-cells. **C)** In T2D, impaired electrophysiologic profiles, assessed as α_probability_ values from three separate classifier models, occur selectively within α-cells enriched in lineage and maturity markers, including *ARX* and *NEUROD1* (see also **Suppl Fig 5**). **D)** The impaired exocytosis in T2D α-cells correlates with expression of *ISL1, NEUROD1, NKX2-2*, or *ARX*. **E)** Exocytosis in ND α-cells at 1 mM glucose was not different in cells enriched in these markers. T2D α-cells with low expression of *ISL1, NEUROD1, NKX2-2*, and *ARX* appear to have normal exocytosis, while cells enriched in these markers exhibit a selective impairment of exocytosis at 1 mM glucose. Data were compared by two-way ANOVA followed by two-stage step-up method for estimation of FDR or Tukey post-test (panel B). *-p<0.05, **-p<0.01, ***-p<0.001 and ****p<0.0001 as indicated.

## Discussion

Glucagon secretion from pancreatic α-cells is under the control of metabolic, paracrine, hormonal, and neuronal signals, to exert its important regulatory role in metabolic homeostasis (El et al., 2020). Much debate has centered around the question of whether glucose-suppression of glucagon secretion is mediated via intrinsic, paracrine, or autonomic mechanisms although it seems likely that α-cells (like β-cells) adeptly integrate multiple signals for precise physiologic control of glucagon. A role for direct glucose-sensing in the α-cell is supported by the impact of cell-specific glucokinase knockout (Basco et al., 2018), the effect of glucokinase activation or knockdown on exocytosis from α-cells (Moede et al., 2020) and *in vivo* glucagon responses (Bahl et al., 2021), and the ability of glucose metabolism to modulate α-cell ATP-sensitive K^+^ channels (Zhang et al., 2013) among other cell-autonomous effects (Gylfe, 2016). Here we show that increasing glucose suppresses exocytosis from human and mouse α-cells stimulated by direct membrane depolarization, consistent with a recent report in human α-cells where exocytosis was measured by live-cell imaging (Omar-Hmeadi et al., 2020).

A role for metabolism in the responses we observe is supported by experiments with the non-metabolizable glucose analog 2-DG, and in the correlation of mitochondrial respiratory complex assembly transcripts with exocytotic responses. Patch-seq analysis highlights components of the electron transport chain complex I which suggests a role for ATP synthesis and/or superoxide production. The study by Omar-Hmeadi et al (2020) also showed a U-shaped response of isolated α-cells to glucose. We find some hints of this in our own data (see Fig 2D) and find that this may depend somewhat on the culture conditions, as we observe a clear glucose-dependent ‘granule refilling’ effect seen following prolonged low-glucose preincubation (**Suppl Fig 6**). This is reminiscent of the glucose-dependent increase in glucagon secretion reported previously from a purified α-cell population (Olsen et al., 2005). Thus, some minimum of glucose metabolism facilitates granule priming or docking to promote a releasable glucagon granule pool, although this may be small given that robust exocytosis persists with 1 mM glucose in the presence of 2-DG but not in the complete absence of glucose. At higher glucose, metabolism acutely suppresses P/Q-type Ca^2+^ channels to limit depolarization induced α-cell exocytosis. Altogether this will ‘tune’ responsiveness to the myriad paracrine, endocrine, and neuronal inputs encountered by the α-cell.

The coupling of glucagon exocytosis to P/Q-type Ca^2+^ channels, which we confirm at low glucose in this study, has been demonstrated previously in α-cells (Marinis et al., 2010; Ramracheya et al., 2010) and is similar to the direct coupling of insulin granule exocytosis in β-cells to the activation of L-type Ca^2+^ channels (Wiser et al., 1999). These and other electrophysiological properties are used to distinguish α- and β-cells from rodents. Most commonly, mouse islet cell types are identified by a combination of cell size (α-cells are smaller) and distinct properties of Na^+^ current inactivation. In a model of genetically induced α-to-μ trans-differentiation in mice, we reported a clear shift in electrophysiological phenotype consistent with the attainment of β-cell properties (Chakravarthy et al., 2017). Intriguingly, following high fat feeding mouse α-cells undergo a negative-shift in Na^+^ current inactivation and convert from P/Q-type to L-type Ca^2+^ channel dependence of exocytosis. While we provide some evidence for impaired α-cell identity, this is not associated with a clear trans-differentiation since β-cell markers are not increased and cells maintain positive immunostaining for glucagon. The shift in Na^+^ channel inactivation towards a ‘β-cell like’ phenotype, which could perhaps be related to changes in membrane composition (Godazgar et al., 2018), correlates with the expression of important α-cell lineage and identity transcription factors, suggesting that the change in electrical phenotype in these cells occurs more readily in cells with higher levels of these markers.

A recent report suggests that a subset of human α-cells, even from donors without diabetes, may exist in an immature state and may suffer a further loss of mature identity in T2D (Avrahami et al., 2020), perhaps related to their greater epigenetic plasticity (Bramswig et al., 2013) and distinct human α-cell states as defined by chromatin accessibility (Chiou et al., 2021). Here we provide evidence that ND α-cells that exhibit what would be considered inappropriately low exocytotic responses are enriched in transcripts and pathways associated with endocrine development (including *FOXO1, PAX6, RFX6* and others) suggesting that α-cells with a lesser degree of maturity (Avrahami et al., 2020) show a concomitant alteration in functional properties. In T2D we see no obvious loss of α-cell transcription factors, or up-regulation of β-cell-defining transcription factors within the α-cell population that would be indicative of trans-differentiation *per se*. We do however find heterogeneity in many lineage markers, as reported previously by us (Camunas-Soler et al., 2020; Drigo et al., 2019) and others (Li et al., 2016). In our dataset, most of these show a small but significant increase in T2D. In order to assess a shift in electrophysiological phenotype of these human α-cells however, we could not use Na^+^ current inactivation as this feature overlaps markedly with measurements from human β-cells. We therefore modified machine learning approaches that we previously used to improve the electrophysiological identification of mouse (Briant et al., 2017) and human (Camunas-Soler et al., 2020) islet cells. We used exocytosis, Na^+^ current, and Ca^2+^ properties as training data for three separate modelling approaches with similar results. Two of these models (Models 2 and 3) excluded cell size, to solely identify shifts in membrane ‘activity’, and all models accurately assigned probability values for ND α- and β-cells irrespective of ambient glucose values, culture times and other important donor and isolation-related parameters. Correlation of α_probability_ values with transcriptomic data therefore emphasized pathways linked to a ‘loss of functional phenotype’ in T2D and suggest that α-cells with the most altered electrical phenotypes in T2D are those with higher expression of pancreatic endocrine lineage and maturity factors. Indeed, we confirmed a selective loss of exocytotic responsiveness at low glucose in T2D α-cells enriched in endocrine progenitor and α-cell maturation markers.

In this study we define an impaired α-cell function phenotype in T2D that is largely restricted to a sub-group of cells expressing higher levels of markers that define not only α-cell maturity, but pancreatic endocrine lineage. At first look we would have expected that high levels of *ARX* should be indicative of mature α-cell function, but this is misleading given a strong overlap of *ARX*^hi^ cells with transcripts more traditionally associated with an endocrine progenitor state such as *NEUROD1, ISL1, FEV* and others. These cells indeed appear more ‘plastic’ in their electrophysiological responses, which may predispose them to an altered electrical phenotype in T2D. While it is tempting to speculate that a ‘de-repression’ of immature gene-sets (Avrahami et al., 2020) drives this altered function, many of the transcripts previously associated with an α-cell ‘toddler’ or ‘juvenile’ phenotype (Arda et al., 2016) are below the level of detection in our data. Nonetheless, aside from highlighting the utility of the patch-seq approach to study heterogenous islet cell populations, our work demonstrates that the function of all α-cells are not equally impacted by disease and that a sub-set of α-cells defined by their maturation state may be key drivers of impaired glucagon responses in T2D.

### Limitations of the study

While we show that glucose metabolism ‘tunes’ the exocytotic responsivity of α-cells, we should be careful when interpreting this. For example, while it is tempting to link impaired exocytosis in T2D α-cells at low glucose to an impaired responsiveness to hypoglycemia *in vivo*, this must be considered in the context of altered responses of paracrine and hormonal signals, which we have not examined here. We studied single isolated α-cells, and although we demonstrate a similar heterogeneity in ARX/MAFB protein expression *in situ*, α-cell function phenotypes are likely to be somewhat different within the intact pancreas. Encouragingly however, we find clear differences between ND and T2D α-cell phenotypes that persist *in vitro* and among the novel findings of this study we also find well-established regulators of a-cell function in situ and in vivo (such as *GIPR* for example). Additionally, our approach to ‘electrophysiological fingerprinting’ is unaffected by time in single-cell culture (up to 3 days), where we might expect phenotypic alterations to become more obvious.

We must also consider that these are not static cell populations we are observing, but rather, they likely represent dynamic cell states. Although we find a selective disruption of function in α-cells defined by high *ARX* and lineage markers, it remains unclear whether exocytotic function would be restored in these cells as they transition from an *ARX*^hi^ to *ARX*^lo^ state. Perhaps this transition itself is impaired in T2D, however we see only modest evidence for an increase in expression of these markers. Finally, we should be careful when directly comparing the rodent and human studies. One clear difference we find is that human T2D α-cells show impaired (low) exocytosis at 1 mM glucose while the mouse HFD α-cells do not (instead, they show altered Na^+^ and exocytosis-Ca^2+^-channel coupling). The exact reasons for this are unclear, perhaps related to obvious differences between the humans and the mouse model (timing, glycemia, age, etc.). Nonetheless, in both the mice and human cells, we find evidence to suggest that α-cell dysfunction is linked to cell maturation state.

## Methods

### Human and mouse islets

In most cases, human islets were from our in-house human islet isolation and distribution program (www.isletcore.ca) (Lyon et al., 2019). Some human islets were provided by the Clinical Islet Transplant Program at the University of Alberta or by Dr. Rita Bottino at the Alleghany Health Network (Pennsylvania, US). Details of donors with no diabetes (ND) or type 2 diabetes (T2D) used in this study are shown in **Supplementary Table 1**. T2D was determined either by reporting of previous clinical diagnosis at the time of organ procurement or, by assessment of HbA1c >6.5% in a few cases that were considered as previously undiagnosed T2D. Human islets and dispersed cells were cultured in DMEM (Gibco 11885) with 10% FBS (Gibco 12483-Canadian origin) and 100 U/ml penicillin/streptomycin (Gibco) at 37°C and 5% CO_2_. Mouse islets were isolated from male C57bl6 mice at 10-12 weeks of age or following 10-12 weeks of high fat diet (60% of calories from fat) starting from 8 weeks of age by collagenase digestion and hand-picking (Smith et al., 2020). Mouse islets and dispersed cells were cultured in RPMI (Gibco 11875) with 10% FBS (Gibco 12483-Canadian origin) and 100 U/ml penicillin/streptomycin (Gibco) at 37°C and 5% CO_2_).

### Patch-clamp recordings

Hand-picked islets were dissociated to single cells using StemPro accutase (Gibco/Fisher, A11105-01). Dispersed islet cells were cultured for 1–3 days, after which media was changed to bath solution containing: 118 mM NaCl, 20 mM TEA, 5.6 mM KCl, 1.2 mM MgCl_2_, 2.6 mM CaCl_2_, 5 mM HEPES, and with glucose as indicated (pH adjusted to 7.4 with NaOH) in a heated chamber (32–35 °C). For whole-cell patch-clamping, fire polished thin wall borosilicate pipettes coated with Sylgard (3–5 MOhm) contained intracellular solution (125 mM Cs-glutamate, 10 mM CsCl, 10 mM NaCl, 1 mM MgCl_2_, 0.05 mM EGTA, 5 mM HEPES, 0.1 mM cAMP and 3 mM MgATP (pH adjusted to 7.15 with CsOH). Electrophysiological measurements were collected using a HEKA EPC10 amplifier and PatchMaster Software (HEKA Instruments Inc., Lambrecht/Pfalz, Germany) within 5 minutes of break-in as described previously (Camunas-Soler et al., 2020). Quality control was assessed stability of the seal (>10 GOhm) and access resistance. Cells were identified by post-hoc immunostaining for insulin with a rabbit anti-insulin primary antibody (cat#SC-9168, Santa Cruz) and goat anti-rabbit Alexa Fluor 488 secondary (cat#A11008, Fisher), and with a guinea pig anti-glucagon primary antibody (cat#4031, Linco) and goat anti-guinea pig Alexa Fluor 594 secondary (cat#A11076, Fisher); or following collection for single-cell RNA sequencing analysis (Camunas-Soler et al., 2020).

For validation of receptor agonist effects on Na^+^ currents (**Fig 5F**), culture media were changed to bath solution containing (in mM): 118 NaCl, 20 TEA, 5.6 KCl, 1.2 MgCl_2_, 5 HEPES, and 5 mM glucose (pH 7.4 with NaOH); and intracellular solution with (in mM): 125 Cs-glutamate, 10 CsCl, 10 NaCl, 1 MgCl_2_, 0.05 EGTA, 5 HEPES (pH 7.15 with CsOH). Compounds used were: lysophosphatidic acid (LPA; Sigma, cat#L7260), α-latrotoxin (Enzo Life Sciences, cat#50-200-9609), Slit guidance ligand 2 (SLIT-2-N; Sigma, cat#SRP3155), prostaglandin E_2_ (PGE2; Tocris, cat#2296), somatostatin (SST; Sigma, cat#S9129), glucose-dependent insulinotropic polypeptide (GIP; Eurogentee, cat#AS-65568), L-glutamic acid (Sigma, cat#G1251), and epinephrine (Sigma, cat#E4375). These were added in bath solution, at the concentrations indicated in the figure legend, and pH was re-adjusted if needed. Na^+^ current was activated by a 50 ms depolarization from -70 to -10 mV.

### Pancreas Patch-seq

Our protocol for pancreas patch-seq is outlined in a recent paper (Camunas-Soler et al., 2020). In brief, following patch-clamp cells were collected using a separate wide-bore collection pipette (0.2-0.5 MOhm) filled with lysis buffer (10% Triton, ribonuclease inhibitor 1:40; Clontech, Cat#2313A), ERCC RNA spike-in mix (1:600000; ThermoFisher, Cat#4456740), 10 mM dNTP, and 100 μM dT (3’-AAGCAGTGGTATCAACGCAGAGTACTTTTTTTTTTTTTTTTTTTTTTTTTTTTTTVN-5’) and then transferred to PCR tubes and stored at -80°C. We generated cDNA and sequencing libraries using an adaption of the SmartSeq-2 protocol for patch-seq plates (Camunas-Soler et al., 2020; Picelli et al., 2014). Libraries were generated from the amplified cDNA by tagmentation with Tn5 and sequenced in the NovaSeq platform (Illumina) using paired-end reads (100 bp) to an average depth of 1 million reads per cell. Sequencing reads were aligned to the human genome (GRCh38 genome with supplementary ERCC sequences) using STAR (Dobin et al., 2013), and gene counts determined using htseq-count (intersection-nonempty) using a GTF annotation with Ensembl 89 release genes (Anders et al., 2015). Gene expression was normalized to counts per million (cpm) after removal of counts corresponding to ERCC spike-ins and transformed to log2 values after addition of a pseudocount. This was followed by QC filtering of the sequenced cells (Camunas-Soler et al., 2020) to remove low-quality cells and potential doublets. Raw sequencing reads are available in the NCBI Gene Expression Omnibus (GEO) and Sequence Read Archive (SRA) under accession numbers GSE124742 and GSE164875. Correlation between transcript expression and electrophysiology was as outlined previously (Camunas-Soler et al., 2020). Visualization as t-distributed stochastic neighbor embedding (tSNE) plots was with the Qlucore Omics Explorer v3.6. Gene-set enrichment analysis (GSEA) was performed with of Z-scores or slopes for transcripts found in >20% of cells using the WEB-based Gene SeT AnaLysis Toolkit (webgestalt.org) and weighted set gene coverage to reduce redundancy in identified terms (Liao et al., 2019).

### Electrophysiological Fingerprint Modelling

Multiple regression was carried out on ND α-cells using Ordinary Least Squared (OLS) Regression from the Statsmodel package in a Python framework. Independent variables included age, sex, body mass index (BMI), HbA1c, cold ischemia time (CIT), culture time, and glucose concentration in the bath solution. Dependent variables included: cell size (pF), total exocytosis (fF/pF), early exocytosis (fF/pF), late exocytosis (fF/pF), peak Na^+^ current (pA/pF), Na^+^ half-Inactivation (mV), early Ca^2+^ current (pA/pF), late Ca^2+^ current (pA/pF), Ca^2+^ charge entry during an initial depolarization (pC/pF), exocytosis normalized to Ca^2+^ charge entry (fF/pC), reversal potential (mV), and α_probability_ from Models 1-3 (see below). Cells lacking data for a dependent variable were only dropped from that specific OLS analysis.

Classification of cell type was conducted using the above electrophysiological measures as dependent variables from ND donors in an Optimizable Ensemble that either included (Model 1) or excluded (Model 2) cell size in MATLAB, or using Extreme Gradient Boosting (XGBoost) in a Python framework that excluded cell size and reversal potential and restricted training data to 32 ND donors with an age of 20-70 years, a BMI of 18.5-30.3 and a pancreas CIT of ≤20 hours. Fine tuning was performed with a pre-determined minimum accuracy of 80% for both α-cell and β-cells. Models were trained on 80% of the α- and β-cells, and 20% were reserved for testing and validation. Confusion matrices were generated using the scikit-learn package in Python. We applied the models across our combined immunostaining and patch-seq database of ND and T2D cells and utilized the classifier’s predicted probability scores to assess fit to α-cell (α_probability_ = 1.0) and β-cell (α_probability_ = 0.0) models.

### Hormone secretion measurements

Groups of 150-175 hand-picked mouse islets were perifused using a BioRep perifusion system (BioRep, Miami). Islets were pre-incubated in perfused KRB buffer containing 140 mM NaCl, 3.6 mM KCl, 2.6 mM CaCl_2_, 0.5 mM NaH_2_PO_4_, 0.5 mM MgSO_4_, 5 mM HEPES, 2 mM NaHCO_3_ and 0.5 mg/ml Essentially fatty acid free BSA (Sigma A6003) for 30 minutes, and then perifused in the same KRB with changes in glucose and KCl as indicated. Samples were collected at 90 -210 sec intervals and stored for assay of glucagon (MSD mouse glucagon kit) and insulin (Alpco Stellux mouse insulin kit) at -20°C.

### Immunostaining, imaging, and single-cell protein analysis

Human paraffin embedded pancreas biopsies were sectioned to 3 μm and immunostained with antibodies against MAFB (Cell Signaling Technologies, cat#41019), ARX (R&D Systems, cat#AF7068) and GCG (Sigma, cat#G2654). Immunofluorescent mages were acquired on a Zeiss Axio Imager M2 widefield microscope with ApoTome. Images containing islet and exocrine cells were processed for nuclei segmentation, individual cell detection and cytosol border inference using Qupath software (version 0.2.3). Parameters for nuclei detection: background radius 20px, median filter 5px, sigma 7px, minimum nuclei area 10px^2^, maximum nuclei area 1000px^2^, threshold of 2, nuclei were split by shape, and cell boundaries were determined by an expansion threshold of 12px with smoothing. Next, the relative expression levels of nuclear ARX, nuclear MAFB and cytosolic GCG levels for each detected cell were exported as .csv files and imported into Cytomap software for spatial analysis (Stoltzfus et al., 2020). In Qlucore Omics Explorer, we performed dimensionality reduction using tSNE of all imaged islet dataset using default parameters with 30 perplexity and 0.5 theta. Data was normalized for each dataset and the relative GCG, ARX and MAFB levels were displayed in the tSNE plot. Next, we created a gate to select the alpha cell population in our dataset and the relative levels of ARX and MAFB were plotted. This information was used to determine the identity of ARX-high and ARX-low cells and the relative expression levels of ARX, MAFB and GCG for each alpha cell subpopulation was plotted.

### Statistical analysis

Data are expressed as mean and standard error (line plots), mean and 10-90 percentile range (box and whisker plots), or as violin plots with median and quartiles indicated. When comparing two groups we used Student’s t-test or the non-parametric Mann-Whitney test to compare ranks. When comparing more than two groups we used one-way or two-way ANOVA followed by either the Tukey post-test or two-stage step-up method for estimation of false discovery rate (FDR); or alternatively the non-parametric Kruskal-Wallis test followed by Dunn’s post-test to correct for multiple comparisons. Statistical tests used are indicated in figure legends. p-values less than 0.05 were considered as significant. For correlation of electrophysiology with transcript expression, Spearman tie-corrected correlations were computed for each gene and significance was tested by bootstrapping (1,000 iterations) as described in our previous work (Camunas-Soler et al., 2020). For ORA and GSEA, false discovery rate (FDR) of reported pathways was <0.05 or <0.1 as indicated in the figures using Bengamini-Hochberg (BH) correction within the WEB-based GEne SeT AnaLysis Toolkit (webgestalt.org).

## Acknowledgements

We thank Dr. Jesper Grud Skat Madsen (University of Southern Denmark) and Dr. Lori Sussel (University of Colorado) for helpful discussion, and Dr. Francis Lynn (University of British Columbia) for critical reading of the draft manuscript. We thank the Human Organ Procurement and Exchange (HOPE) program and Trillium Gift of Life Network (TGLN) for their work in procuring human donor pancreas for research. We also thank Dr. Rita Bottino (Allegheny Health Network) and Drs. James Shapiro and Tatsuya Kin (University of Alberta Clinical Islet Program) for contributing some islet preparations for this study. Finally, we especially thank the organ donors and their families for their kind gift in support of diabetes research.

TdS was supported by the Alberta Diabetes Institute (AD)-Helmholtz Diabetes Research School for Diabetes, the Alberta Innovates Scholarship in Data-Enabled Innovation, and the Sir Fredrik Banting and Dr. Charles Best Canada Graduate Scholarship. This work was funded by grants to PEM from the Canadian Institutes of Health Research (CIHR: 148451), from the JDRF to PEM, LB and PR (SRA-2019-698-S-B), and from the National Institutes of Health to EMW (F32 DK109577) and to PEM, RAD, SK (U01DK120447). PEM holds the Canada Research Chair in Islet Biology.

## Author Contributions and Guarantor

XD, JC-S, AB, AFS, TdS, LB, AN, RAD, EW and RJ collected and analyzed data. JL, NS, AB and JEMF isolated human islets. SK, SQ, PR, RS and PEM conceived the study, designed experiments, and analyzed data. PEM wrote the initial manuscript draft and all authors contributed to editing the final version. PEM acts as the guarantor of this work and is responsible for data access.

