## Supplemental Figures for "Heterogenous impairment of α-cell function in type 2 diabetes is linked to cell maturation state"

#### Supplementary Figure 1

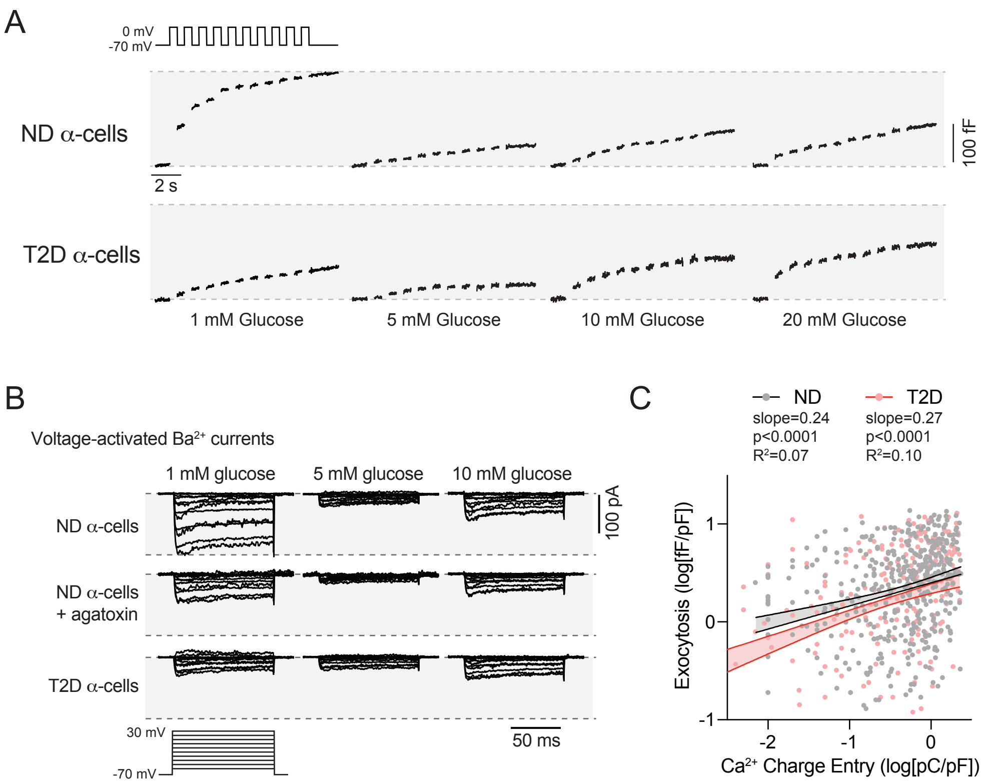


**A)** Exocytosis was elicited from human alpha-cells with a series of membrane depolarizations from -70 to 0 mV and measured as increases in membrane capacitance. Representative traces are shown (see main Fig 1 for summarized data). In alpha-cells from donors with no diabetes (ND), exocytosis responses were greatest with 1 mM glucose in the bath and decreased at higher glucose levels. In alpha-cells from donors with type 2 diabetes (T2D) exocytosis was persistently low at 1 mM glucose and, if anything, was slightly facilitated by increasing glucose. **B)** Voltage-dependent $Ca^{2+}$ channel activity, measured using $Ba^{2+}$ as a charge carrier, was suppressed with increasing glucose in ND alpha-cells. Blockade with agatoxin (100 nM) shows that most of the increased current at 1 mM glucose is mediated by P/Q-type $Ca^{2+}$ channels. This current appears to be missing in T2D alpha-cells. **C)** Comparing the initial exocytotic response with the $Ca^{2+}$ influx during an initial (1, 5, 10 mM glucose combined) shows a significant relationship between $Ca^{2+}$ entry and alpha-cell exocytosis that is unaffected by T2D, suggesting the reduced exocytosis seen in T2D results from the reduced $Ca^{2+}$ influx.

### Supplementary Figure 2

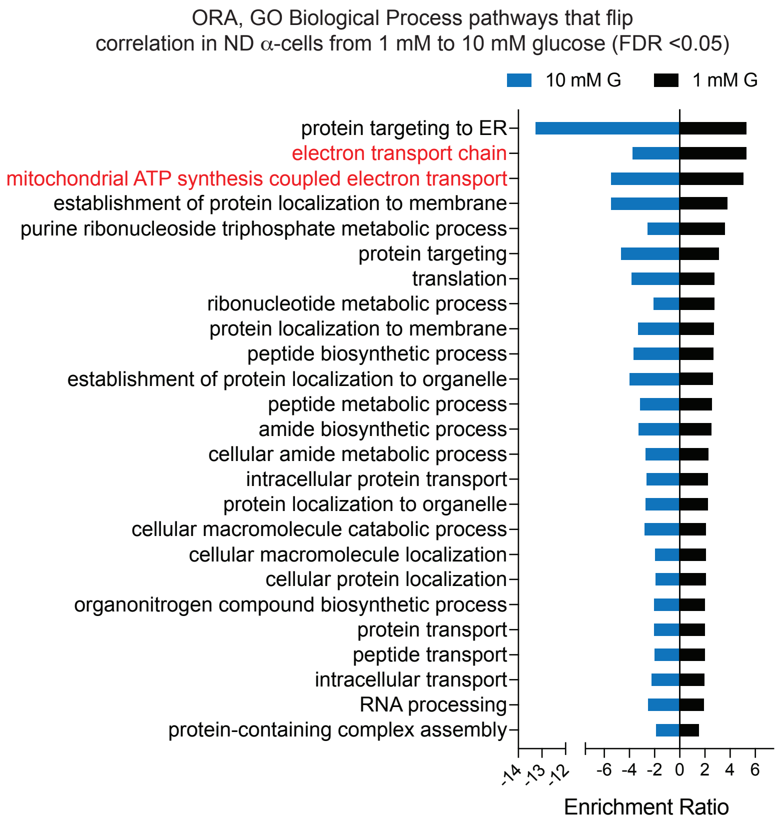


In alpha-cells from donors with no diabetes (ND), GO Biological Process pathways identified by over-representation analysis (ORA) of transcripts with significant ($p<0.05$) positive correlation with total exocytosis at 1 mM glucose (black) or negative correlation with exocytosis at 10 mM glucose (blue) listed in Suppl Table 2. ORA was performed with transcripts expressed in >20% of alpha-cells using the WEB-based GEne SeT AnaLysis Tool (webgestalt.org).

### Supplementary Figure 3

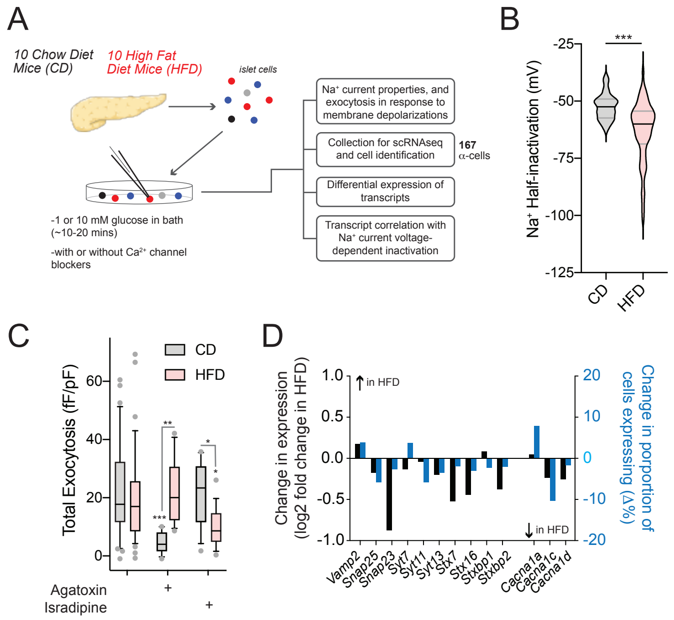


**A)** We analyzed 167 $\alpha$-cells from 10 male C57/bl6 mice fed chow diet (CD) and 10 male C57/bl6 mice fed high fat diet (HFD) by patch-seq. **B)** We confirm in this patch-seq dataset that mouse $\alpha$-cell $Na^+$ current inactivation shifts to more negative voltages following HFD (D; n= 50, 65 cells). **C)** We confirm that mouse $\alpha$-cell exocytosis at 1 mM glucose shifts from P/Q-channel to L-type $Ca^{2+}$ channel dependence in HFD $\alpha$-cells (n=32,37,18,12,12,16 cells). **D)** This is associated with down-regulation of the expression (*black*) and percent of cells expressing (*blue*) some exocytosis-related transcripts. Error bars in box-and-whisker plots show the 10-90% percentile range. Data were compared by two-way ANOVA followed by Tukey post-test or the Student's T-test. *-p<0.05; **-p<0.01; and ***-p<0.001 compared with the 1 mM glucose control or as indicated.

#### Supplementary Figure 4

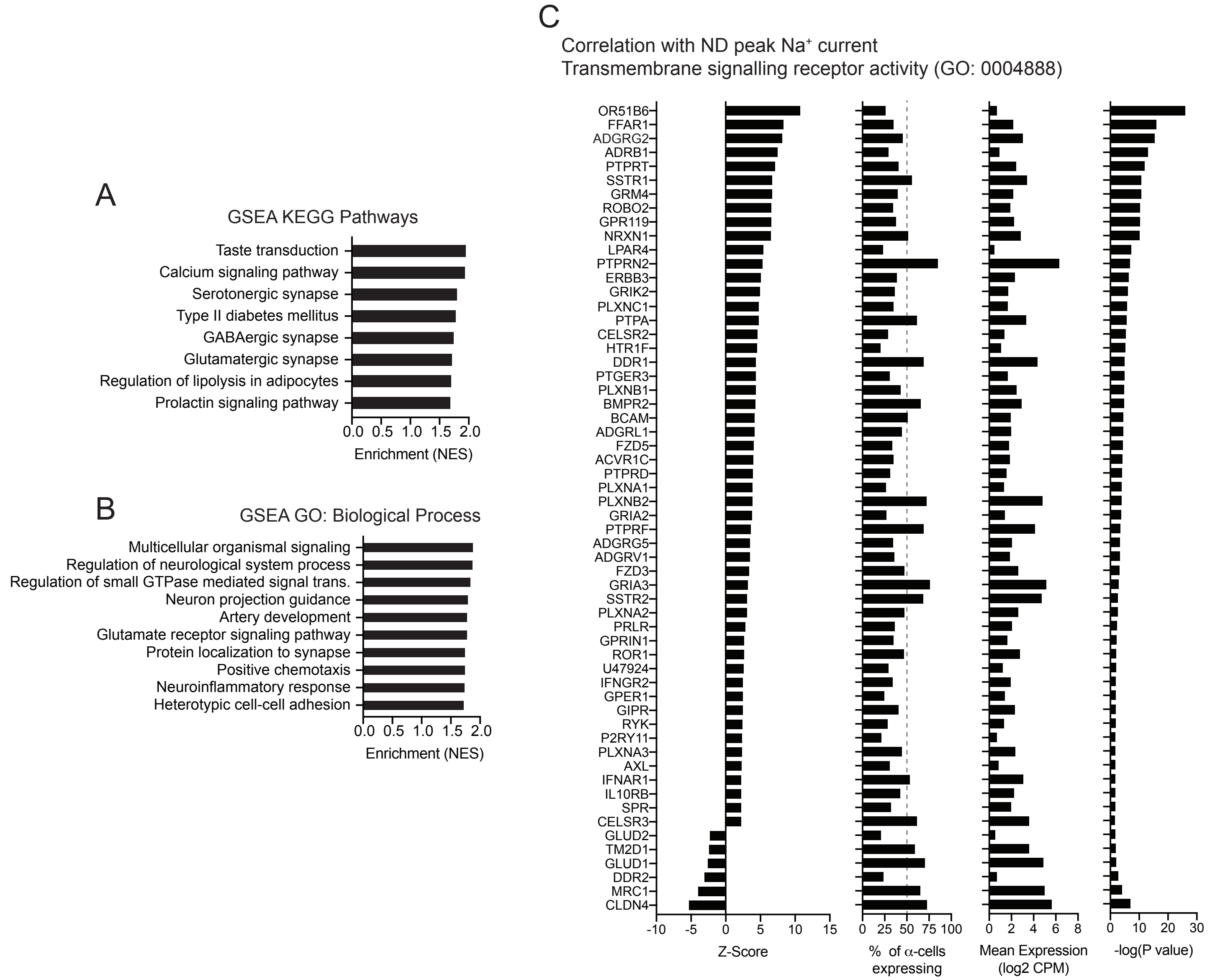


**A-B)** Gene set enrichment analysis (GSEA) using peak $Na^+$ current correlation Z-scores of transcripts expressed in at least 20% of alpha-cells as weighthing identifies several KEGG Pathways (A) and GO: Biological Process Pathways (B), including numerous involved in receptor and neurotransmitter signalling. False discovery rate (FDR) <0.05 for all pathways shown.

**C)** In alpha-cells from donors with no diabetes (ND), transmembrane signalling receptors (GO: 0004888) with significant ($p<0.05$) correlation to sodium current peak amplitude.

### Supplementary Figure 5

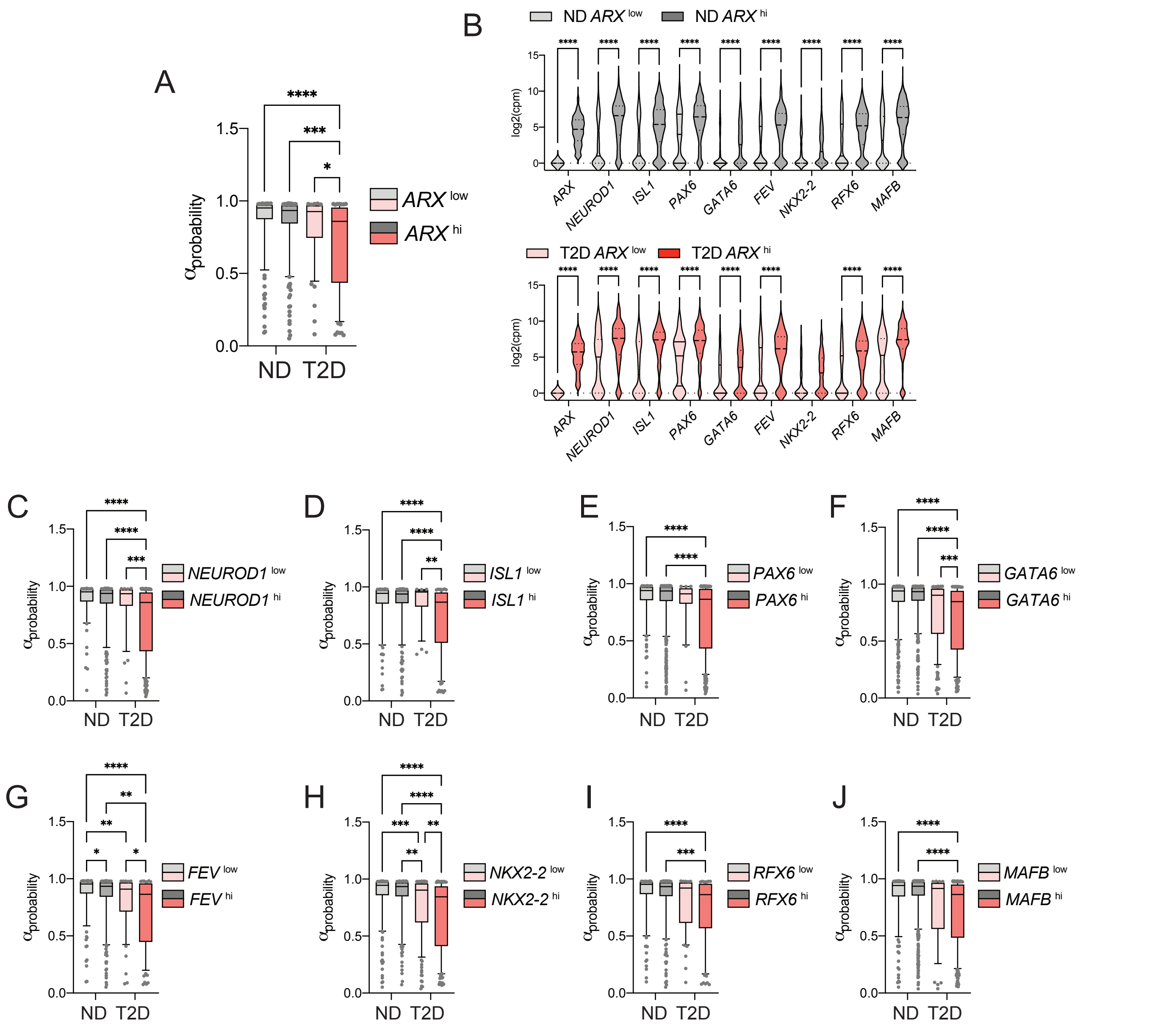


**A)** Human alpha-cells were separated based on the low or high expression of ARX (*ARX*lo; *ARX*hi). In type 2 diabetes (T2D; *pink/red*) a loss of electrophysiological identity, assessed using $\alpha_{probability}$ values from our Model 3 (XGBoost with training restrictions to donor age, BMI and cold ischemic time), is observed selectively in *ARX*hi cells. **B)** The *ARX*hi alpha-cells also express higher levels of many progenitor and linage markers both in cells from donors with no diabetes (ND; *light/dark grey*) and with T2D. **C)** Separating alpha-cells based on low and high expression of each of these markers we similarly find a selective loss of electrophysiological phenotype in the ‘high’ marker expressing cells (again, using Model 3 $\alpha_{probability}$ values).

### Supplementary Figure 6

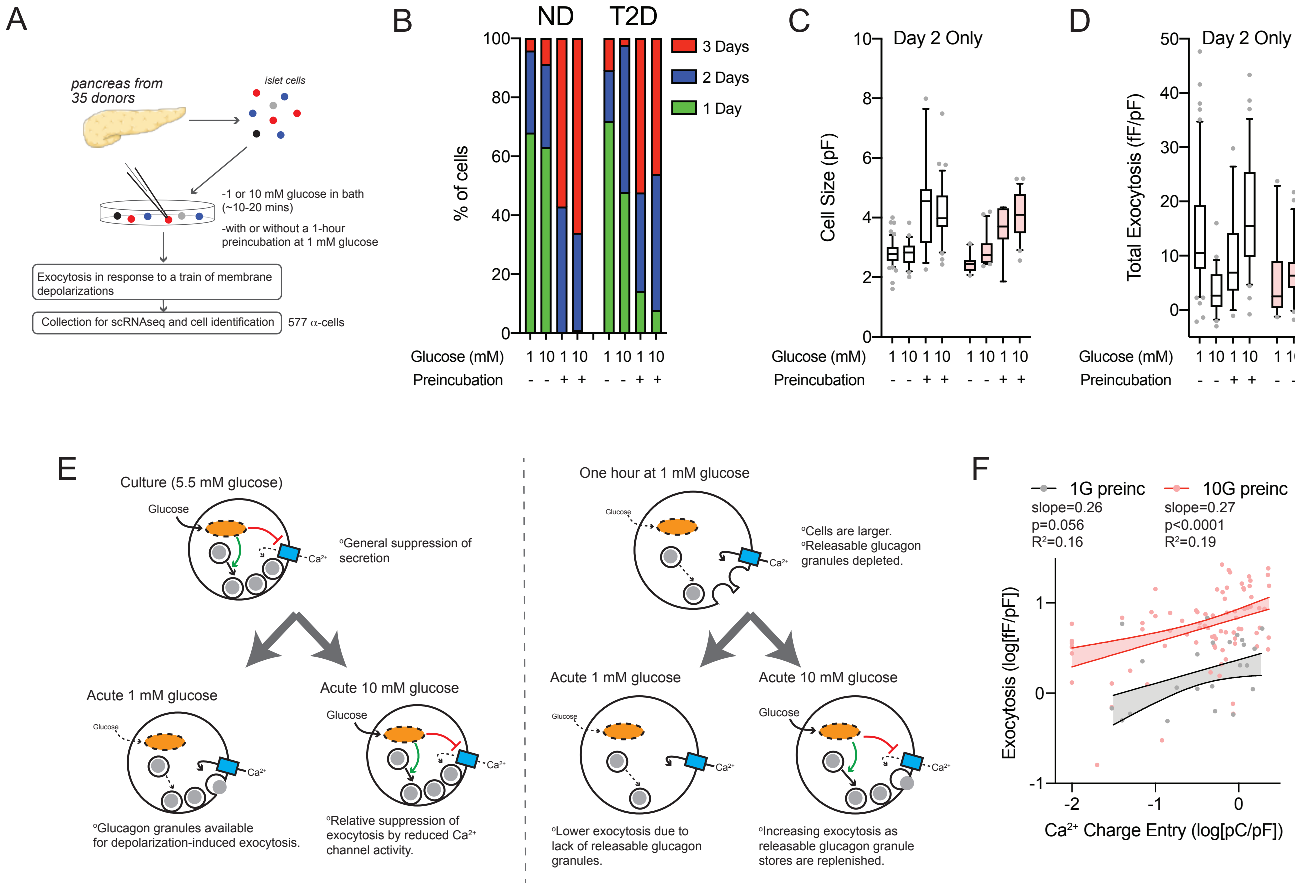


Comparison of human alpha-cell size and exocytosis upon acute glucose change (following culture at 5.5 mM glucose) or following a 1-hour preincubation at 1 mM glucose. **A**) We collected cells either with or without 1-hr preincubation at 1 mM glucose. **B**) The non-preincubated cells were mostly patch-clamped on days 1 and 2, while the 1 mM glucose preincubated cells were mostly patch-clamped on days 2 and 3 following dispersion to single cells. To directly compare the effect of low glucose preincubation, here we looked at cells from day 2 only. **C, D**) We see that low-glucose preincubation increases cells size (C) and under this condition acute stimulation with glucose now facilitates exocytosis (D). The scheme in panel **E** representes what we believe is happening in these experiments. Alpha-cell activity during the 1-hour preincubation at low glucose may exhaust the pool of releasable glucagon granules (the cells being bigger as a result of continued glucagon granule fusion with the plasma membrane). The poor exocytotic response at 1 mM glucose is a consequence of this depletion, and the increasing exocytosis upon acute switching to 10 mM glucose is reflective of the replenishment of the releasable granule pool. **F**) Consistent with this, in 1 mM glucose preincubated cells the slope of the relationship between $Ca^{2+}$ entry and exocytosis is not different at 1 mM glucose or after acute (~10 minute) switching to 10 mM glucose. But the overall amplitude of the exocytotic response is shifted upward by 10 mM glucose, reflective of replenishment the exocytotic pool.
